# Binding of Filamentous Actin to CaMKII as a Potential Mechanism for the Regulation of Bidirectional Synaptic Plasticity by *β*CaMKII in Cerebellar Purkinje Cells

**DOI:** 10.1101/521542

**Authors:** Thiago M. Pinto, Maria J. Schilstra, Antonio C. Roque, Volker Steuber

## Abstract

Calcium-calmodulin dependent protein kinase II (CaMKII) regulates many forms of synaptic plasticity, but little is known about its functional role during plasticity induction in the cerebellum. Experiments have indicated that the *β* isoform of CaMKII controls the bidirectional inversion of plasticity at parallel fibre (PF)-Purkinje cell (PC) synapses in cerebellar cortex. Because the cellular events that underlie these experimental findings are still poorly understood, we developed a simple computational model to investigate how *β*CaMKII regulates the direction of plasticity in cerebellar PCs. We present the first model of AMPA receptor phosphorylation that simulates the induction of long-term depression (LTD) and potentiation (LTP) at the PF-PC synapse. Our simulation results suggest that the balance of CaMKII-mediated phosphorylation and protein phosphatase 2B (PP2B)-mediated dephosphorylation of AMPA receptors can determine whether LTD or LTP occurs in cerebellar PCs. The model replicates experimental observations that indicate that *β*CaMKII controls the direction of plasticity at PF-PC synapses, and demonstrates that the binding of filamentous actin to CaMKII can enable the *β* isoform of the kinase to regulate bidirectional plasticity at these synapses.

**Author Summary:** Many molecules and the complex interactions between them are involved in synaptic plasticity in the cerebellum. However, the exact relationship between cerebellar plasticity and the different signalling cascades remains unclear. Calcium-calmodulin dependent protein kinase II (CaMKII) is an important component of the signalling network that is responsible for plasticity in cerebellar Purkinje cells (PCs). The CaMKII holoenzyme contains different isoforms such as *α*CaMKII and *β*CaMKII. Experiments with *Camk2b* knockout mice that lack the *β* isoform of CaMKII demonstrated that *β*CaMKII regulates the direction of plasticity at parallel fibre (PF)-PC synapses. Stimulation protocols that induce long-term depression in wild-type mice, which contain both *α* and *β*CaMKII, lead to long-term potentiation in knockout mice without *β*CaMKII, and vice versa. We developed a kinetic simulation of the phosphorylation and dephosphorylation of AMPA receptors by CaMKII and protein phosphatase 2B to investigate how *β*CaMKII can control bidirectional synaptic plasticity in cerebellar PCs. Our simulation results demonstrate that the binding of filamentous actin to *β*CaMKII can contribute to the regulation of bidirectional plasticity at PF-PC synapses. Our computational model of intracellular signalling significantly advances the understanding of the mechanisms of synaptic plasticity induction in the cerebellum.

## Introduction

The structure and neural circuitry of the cerebellum are well understood. The precision of the cerebellar anatomy has instigated the development of many theories that attempt to unravel cerebellar function. However, although it is known that the cerebellum contributes to motor learning and cognition, there is still no general agreement about its exact functional role [1–4].

Synaptic plasticity is an activity-dependent change in the strength of synaptic connections between pre and postsynaptic neurons. In cerebellar cortex, modifications in the strength of synaptic connections between parallel fibres (PFs) and Purkinje cells (PCs) such as long-term depression (LTD) and long-term potentiation (LTP) are thought to contribute to cerebellar learning [5–7].

PF LTD (often called cerebellar LTD) is a process in which the strength of the PF-PC synapse is depressed by large increases in the postsynaptic calcium concentration in response to the coincident activation of PF and climbing fibre (CF) input onto the PC [8–13]. In addition to undergoing LTD, PF synapses can also exhibit LTP. The strengthening of excitatory synapses between PFs and PCs by PF LTP is mediated by smaller calcium concentration increases that can result from the activation of PFs without any coincident CF input to the PC. LTP is necessary to balance LTD at cerebellar PF-PC synapses in order to prevent saturation and to allow reversal of motor learning [14]. A large number of experimental [10, 15–21] and several computer simulation studies [11, 13, 22–27] have explored the biochemical pathways involved in PF LTD, but less is known about the mechanisms underlying PF LTP.

Calcium-calmodulin dependent protein kinase II (CaMKII), which is one of the most abundant proteins in the brain, is a multifunctional enzyme that phosphorylates a wide range of substrates [28–31]. CaMKII is a critical mediator of the calcium signalling systems that underlie the induction of synaptic plasticity. Although significant progress has been made in understanding the role of CaMKII in synaptic plasticity in other brain areas, little is known about its functional role during plasticity induction in the cerebellum.

Two CaMKII isoforms, *α*CaMKII and *β*CaMKII, have been shown to mediate synaptic plasticity in the cerebellum, and therefore to be essential for cerebellar learning and memory formation [29, 31–33]. Although *β*CaMKII is the predominant isoform of CaMKII in the cerebellum, the exact role of *β*CaMKII in cerebellar learning has yet to be established.

Experiments with *Camk2b* knockout mice, which have been genetically modified to prevent the expression of the gene encoding the *β* isoform of CaMKII, have addressed the role of *β*CaMKII in plasticity in cerebellar PCs. These studies demonstrated that the *β*CaMKII isoform regulates the direction of cerebellar plasticity at PF-PC synapses [33]. Stimulation protocols that induce LTD in wild-type mice result in LTP in knockout mice that lack *β*CaMKII, and vice versa (Figure 1). However, the underlying mechanism that may explain these experimental findings is not clear. Van Woerden et al [33] suggested that a biochemical difference between the *α*CaMKII and *β*CaMKII isoforms could underlie the switch of the direction of synaptic plasticity. The *β*CaMKII, but not *α*CaMKII, isoforms can bind to F-actin [34], which could result in sequestering of the CaMKII complex to F-actin, making it unavailable for AMPA receptor phosphorylation.

**Figure 1.**
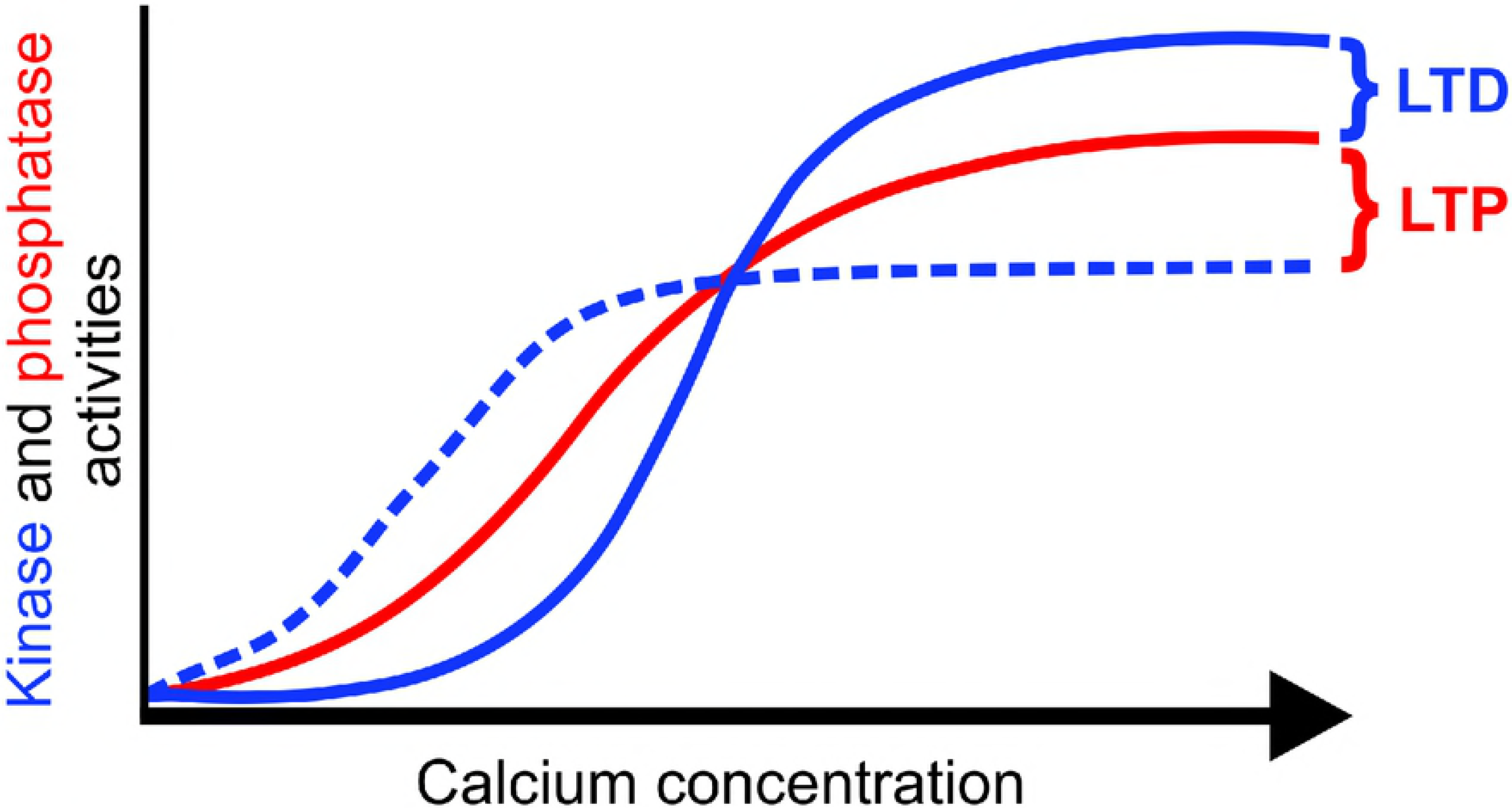
Schematic representation of bidirectional plasticity at PF-PC synapses. The experimental results obtained by Van Woerden and collaborators [33] are schematically represented in this figure. The scheme illustrates how changes in the CaMKII-driven pathway could evoke different activity levels of calcium-dependent kinase (blue), resulting in the inversion of plasticity for wild-type and *Camk2b* knockout mice (solid and dashed lines, respectively). LTD is generated when the kinase activity (blue) surpasses the phosphatase activity (red), whereas the opposite case induces LTP.

Because experimental observations by themselves have not been able to explain how CaMKII mediates plasticity in PCs, a new dynamic model of cerebellar LTD and LTP that includes CaMKII has the potential to enhance our understanding of cerebellar plasticity. To investigate the proposed function of *β*CaMKII in cerebellar plasticity, we developed a mathematical model of the biochemical pathways underlying the calcium-dependent phosphorylation and dephosphorylation of AMPA receptors at PF-PC synapses. Computer simulations of our model were used to explore how *β*CaMKII can mediate the observed reversal of plasticity in cerebellar PCs. Our results indicate that the binding of F-actin to CaMKII can indeed enable the *β*CaMKII isoform to control the direction of plasticity at PF-PC synapses, as proposed by Van Woerden et al [33]. Moreover, our model predicts that the sign inversion of synaptic plasticity is based on two additional mechanisms - a reduction of the overall level of CaMKII in the *Camk2b* knockout mice, and a resulting increase in phosphatase activity in these mice. We present the first data-driven model of intracellular signalling pathways in cerebellar PCs that includes CaMKII and that is able to replicate the induction of LTD and LTP at this important cerebellar synapse.

## Methods

To understand the role of *β*CaMKII in cerebellar LTD and LTP at PF-PC synapses, we developed a mathematical model of the phosphorylation and dephosphorylation of AMPA receptors by CaMKII and protein phosphatase 2B (PP2B). The simple model of AMPA receptor phosphorylation consists of six reactions: calcium-calmodulin (Ca_4_CaM)-dependent activation of CaMKII, CaMKII binding to F-actin, binding of calcium to calmodulin (CaM) to form Ca_4_CaM, PP2B activation by Ca_4_CaM, and AMPA receptor phosphorylation and dephosphorylation by, respectively, CaMKII and PP2B (Figure 2).

**Figure 2.**
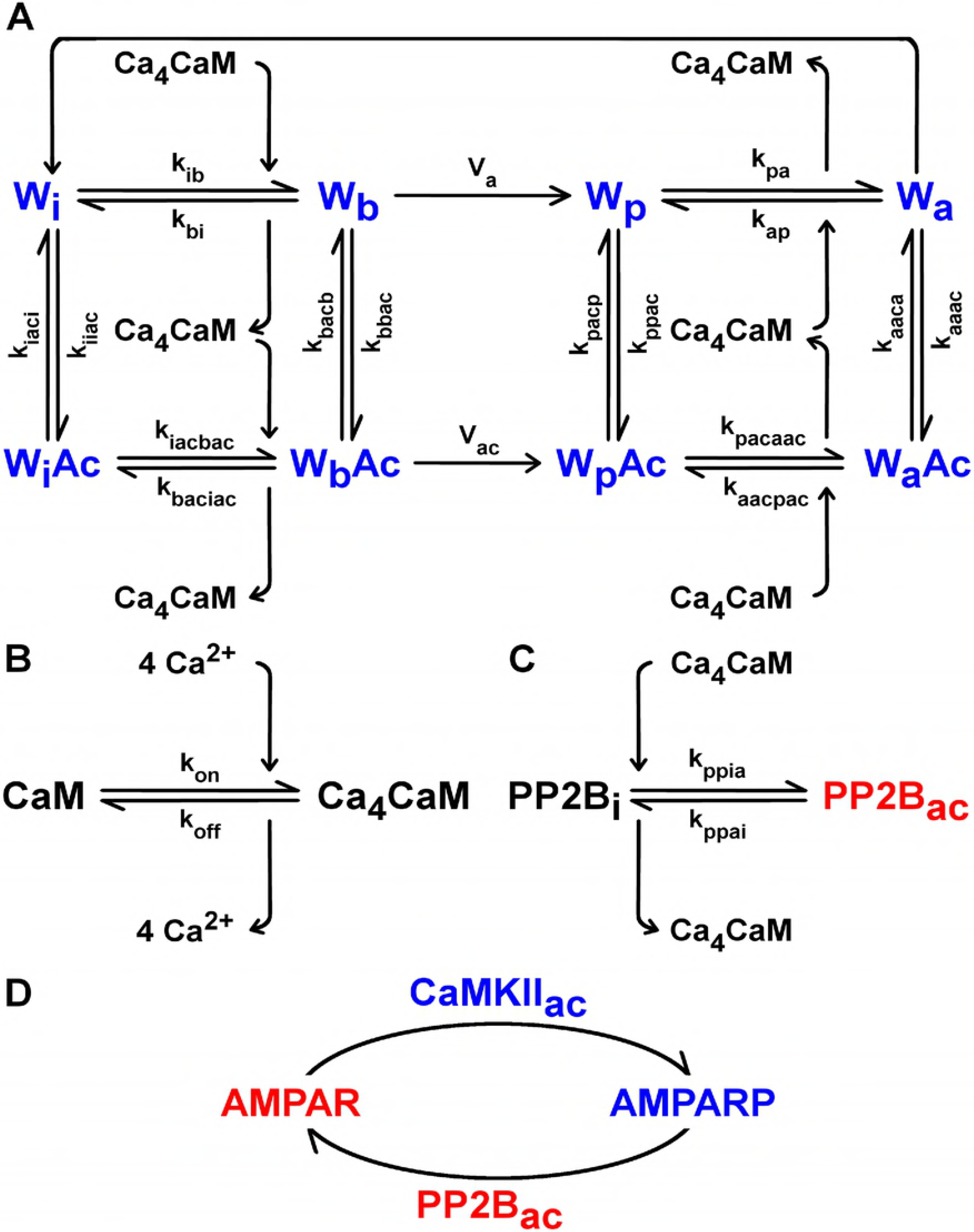
Model of bidirectional plasticity at PF-PC synapses. **A.** CaMKII activation by Ca_4_CaM, and its binding to F-actin (Ac). CaMKII subunits for simulations of *Camk2b* knockout mice are in one of four states: W_i_, W_b_, W_p_, and W_a_, where the subscripts i, b, p, and a refer to the respective subunit states: inactive, bound to Ca_4_CaM, phosphorylated and bound to Ca_4_CaM, and autonomous: phosphorylated, but dissociated from Ca_4_CaM. The kinase can also bind to Ac and be in the W_i_Ac, W_b_Ac, W_p_Ac, and W_a_Ac subunit states, but only in simulations of wild-type mice. Here, we allow the CaMKII subunits in the W_a_ form to switch to the W_i_ state. CaMKII is therefore gradually inactivated after calcium stimulation stops. The kinetic constants of the reversible Ca_4_CaM binding reactions are k_ib_, k_bi_, k_pa_, k_ap_, k_iacbac_, k_baciac_, k_pacaac_ and k_aacpac_, whereas k_iiac_, k_iaci_, k_bbac_, k_bacb_, k_ppac_, k_pacp_, k_aaac_ and k_aaca_ denote the speed of the reversible Ac binding reactions. The rates of the irreversible phosphorylation of W_b_ and W_b_Ac are V_a_ and V_ac_, respectively. **B.** Binding of four calcium ions (4 Ca^2+^) to CaM to form Ca_4_CaM. k_on_ and k_off_ are the rate constants of the reversible 4 Ca^2+^ binding reaction. **C.** PP2B activation by Ca_4_CaM. PP2B_i_ and PP2B_ac_ are the inactive and active forms of PP2B. The respective rate constants of PP2B activation and inactivation are k_ppia_ and k_ppai_. **D.** AMPA receptor phosphorylation and dephosphorylation by active CaMKII (CaMKII_ac_ = W_b_ + W_p_ + W_a_) and PP2B_ac_.

To simulate the mechanism of CaMKII autophosphorylation, we adopt a simplified version of the commonly used model of CaMKII activation by Ca_4_CaM developed by Dupont and collaborators [35]. Because Van Woerden and collaborators suggested that the binding of F-actin to CaMKII could underlie the switch of direction of plasticity [33], we incorporated the binding of F-actin to CaMKII into this model to simulate the plasticity induction in wild-type mice, whereas the F-actin binding was omitted in the knockout mice that lack *β*CaMKII.

In the simple model of Ca_4_CaM-mediated CaMKII autophosphorylation considered here, all CaMKII subunits of the simulated *Camk2b* knockout mice are in one of four states. Following the terminology and notational convention used by Dupont et al [35], the four states are referred to as W_i_, W_b_, W_p_, and W_a_, where the subscripts i, b, p, and a stand for *inactive*, *bound*, *phosphorylated*, and *autonomous*, respectively. W_i_ and W_b_ are unphosphorylated, and W_p_ and W_a_ are phosphorylated states, while subunits in the W_b_ and W_p_ states have Ca_4_CaM bound, and W_i_ and W_a_ have not. The CaMKII subunits of wild-type mice can also be in states bound to F-actin (Ac): W_i_Ac, W_b_Ac, W_p_Ac, and W_a_Ac.

The binding of four calcium ions to CaM to form Ca_4_CaM is included in the model. Furthermore, the Ca_4_CaM complex not only activates CaMKII, but is also responsible for the activation of PP2B. Because binding of *β*CaMKII to F-actin is thought to result in sequestration of the kinase holoenzyme [36], W_b_Ac, W_p_Ac and W_a_Ac are unavailable for AMPA receptor phosphorylation. In the model presented here, phosphorylation of AMPA receptors is therefore mediated by the active CaMKII subunits that are not bound to F-actin (CaMKII_ac_ = W_b_ + W_p_ + W_a_). All of the active PP2B contributes to the receptor dephosphorylation.

Although the cerebellum contains four times as much *β*CaMKII as *α*CaMKII [29], the actual *α*:*β*CaMKII ratio is 1:1 in PCs [32, 33]. Therefore, the CaMKII concentration for *Camk2b* knockout mice is set to half of the kinase concentration for wild-type mice in our model. Moreover, in accordance with experimental observations by Brocke et al that suggest that CaMKII isoforms have different affinities for Ca_4_CaM [37], *β*CaMKII has a greater affinity for Ca_4_CaM than *α*CaMKII in our computer simulations.

All biochemical reactions in our model are represented by coupled ordinary differential equations (ODEs). The full kinetic model with its associated ODEs and the parameter values adopted in all simulations are given in the Supporting Information. All of the following results were obtained by numerically integrating these equations in XPPAUT (X-Windows Phase Plane plus Auto), a numerical tool for simulating dynamical systems, using the CVODE method with relative and absolute error tolerances of 10^−10^.

## Results

Using simulations of our mathematical model, we first investigated the dependence of kinase and phosphatase activities on the postsynaptic intracellular calcium concentration for wild-type mice and *Camk2b* knockout mice. We varied the amplitude of calcium pulses, which are used as input to our model, and calculated the average concentrations of all biochemical compounds during the simulated period for each level of calcium concentration. The goal of these simulations was to compare the predictions of our computational model with the proposed explanation of the experimental findings by Van Woerden and collaborators [33] that is illustrated in schematic form in Figure 1.

Figure 3A shows the resulting average concentrations of CaMKII_ac_ and PP2B_ac_ for wild-type mice and *Camk2b* knockout mice as we increase the average concentration of calcium during stimulation period. To create similar conditions as in a typical cerebellar plasticity induction protocol, which was also used by Van Woerden and colleagues [33], the input to the model consisted of calcium pulses applied at a rate of 1 Hz for 300 s. The figure compares the resulting kinase and phosphatase concentrations averaged over the 300 s of calcium application with concentrations averaged over a longer time period of 6000 s in order to explore the continued behaviour of the system after the offset of calcium application.

**Figure 3.**
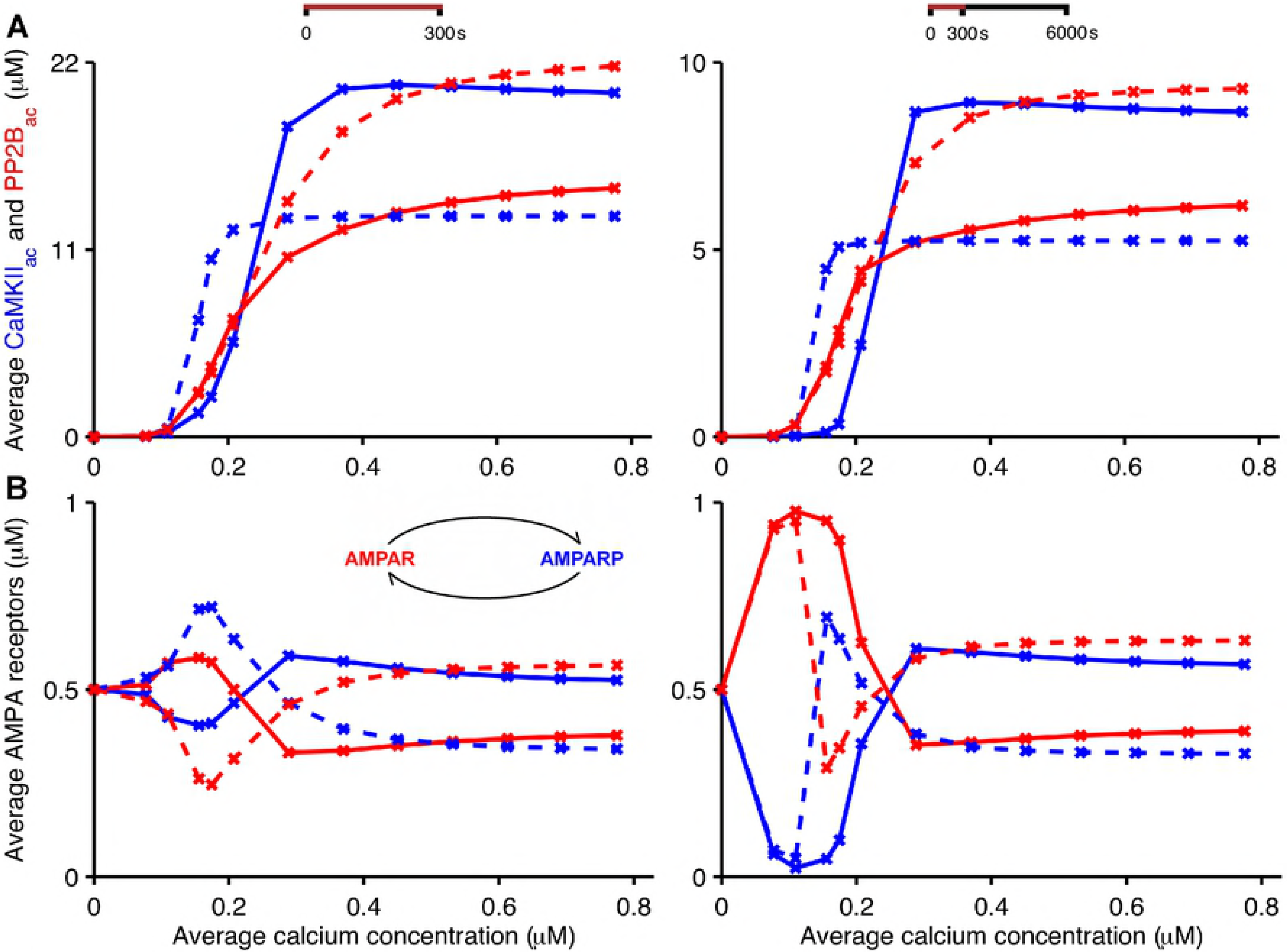
Bidirectional plasticity at PF-PC synapses: kinase and phosphatase activities at different calcium concentrations. The figure shows simulation results that support the suggested explanation of the experimental observations in [33] (Figure 1). Low calcium concentrations that induce LTP in wild-type mice (solid) lead to LTD in *Camk2b* knockout mice (dashed), and vice versa. In the model, the F-actin binding to CaMKII occurs in wild-type mice, whereas in knockout mice the lack of *β*CaMKII prevents the binding of CaMKII to F-actin. **A.** Average concentrations of CaMKII_ac_ (blue) and PP2B_ac_ (red) as a function of average concentrations of pulsed calcium at 1 Hz for 300 s. **B.** Average concentrations of unphosphorylated and phosphorylated AMPA receptors (red and blue, respectively) in response to the same stimulation protocols. In the left panels, all simulations last 300 s, that is, the panel shows average concentrations of CaMKII_ac_, PP2B_ac_ and AMPA receptors in the presence of calcium signals applied at 1 Hz for 300 s. The right panels show the average concentrations of these compounds during simulations that last 6000 s, that is, continue for 5700 s after the offset of the 300 s stimulation.

In both cases, when concentrations were measured during and after the calcium stimulus, our simulation results confirm the explanation of the CaMKII-dependent switch of bidirectional plasticity that was originally put forward by Van Woerden et al [33]. Low calcium concentrations, which would be expected to result from PF input alone, evoke larger average concentrations of PP2B_ac_ than CaMKII_ac_ in simulated wild-type mice. In contrast, for *Camk2b* knockout mice, in which the lack of *β*CaMKII prevents the binding of the CaMKII holoenzyme to F-actin, this relationship is reversed, with low calcium concentrations activating CaMKII more strongly than PP2B. Furthermore, high calcium concentrations, in the range of those expected to be caused by coincident PF and CF input, lead to higher concentrations of CaMKII_ac_ than PP2B_ac_ in wild-type mice, but larger phosphatase than kinase activities in *Camk2b* knockout mice (Figure 3A).

However, in spite of the similarity of our simulation results with the mechanism predicted by Van Woerden and collaborators [33], a comparison of Figures 1 and 3A also reveals a subtle difference. In the schematic originally proposed by Van Woerden et al (Figure 1), the CaMKII-dependent synaptic switch was assumed to be entirely mediated by a change in CaMKII activity due to the genetic *Camk2b* knockout, with low calcium concentrations resulting in increased CaMKII activity in knockout mice, while high calcium concentrations were proposed to lead to reduced CaMKII activity compared to wild-type mice. Our simulation results (Figure 3A) indicate that the proposed reduction in CaMKII activity for larger calcium concentrations is augmented by a coincident increase in PP2B activity. These results predict that an increase in the PP2B_ac_ concentration acts in tandem with a decrease in the CaMKII_ac_ concentration to convert LTD to LTP for large calcium concentrations, or coincident PF and CF input, in the *Camk2b* knockout mice.

The effect of CaMKII and PP2B on cerebellar synaptic plasticity is mediated by the phosphorylation and dephosphorylation of AMPA receptors, respectively, with AMPA receptor phosphorylation leading to the internalization and removal of AMPA receptors from the postsynaptic membrane and the induction of LTD. Thus, we also investigated the effect of the simulated *Camk2b* knockout on AMPA receptor phosphorylation for different calcium concentrations. Figure 3B shows the resulting average concentrations of phosphorylated and unphosphorylated AMPA receptors as the amount of postsynaptic intracellular calcium increases for wild-type and *Camk2b* knockout mice. For simplicity and in the absence of any other knowledge, we assume that the initial concentrations of phosphorylated and unphosphorylated AMPA receptors at a calcium concentration of zero are equal, that is, half of the AMPA receptors are unphosphorylated and the other half are phosphorylated. As in Figure 3A, we compare average concentrations of AMPA receptors during the 300 s calcium application protocol (Figure 3B, left panel) with those averaged over 6000 s (Figure 3B, right panel), lasting for 5700 s after the offset of the calcium stimulus. When measured during the 300 s period of calcium application, the concentration of unphosphorylated AMPA receptors in wild-type mice initially increases with an increased calcium concentration, peaks at an average calcium concentration of about 0.2 *µ*M, and drops off below its initial value for higher calcium concentrations. As expected from the simulated activation of CaMKII and PP2B (Figure 3A), *Camk2b* knockout mice show the opposite behavior: the average concentration of unphosphorylated AMPA receptors during the stimulation period first decreases as the calcium concentration is raised, reaches a minimum where the unphosphorylated AMPA receptors in wild-type mice are at their maximum, and rises above its initial value for higher calcium concentrations. A similar behaviour is observed when the concentration of receptors is averaged over 6000 s, but there is a noticeable difference at very low calcium concentrations (below 0.1 *µ*M) where the levels of unphosphorylated and phosphorylated AMPA receptors are similar for both types of mice. For these very low calcium concentrations, CaMKII is only weakly activated and the concentration of CaMKII_ac_ rapidly drops after the calcium stimulation stops. In this case, the CaMKII_ac_ decay is much faster than the deactivation of PP2B, favouring the dephosphorylation of AMPA receptors (compare Figure 6A for wild-type mice). In other respects, however, the results for 6000 s replicate those for 300 s, with increased (decreased) concentrations of unphosphorylated AMPA receptors for wild-type (knockout) mice at low calcium concentrations at around 0.2 *µ*M, and decreased (increased) levels of unphosphorylated AMPA receptors for wild-type (knockout) mice for higher calcium concentrations (above 0.3 *µ*M). Given that the level of unphosphorylated (and not internalized) AMPA receptors determines the strength of the PF-PC synapses, our simulations therefore replicate the experimental results by Van Woerden et al [33]: in wild-type mice, LTP occurs at low calcium concentrations and LTD at high calcium concentrations, while knockout mice that lack *β*CaMKII exhibit LTD at low calcium concentrations and LTP at high calcium concentrations.

To examine in more detail how the concentrations of the different biochemical species evolve over time, we analysed the temporal evolution of kinase and phosphatase activities and AMPA receptor concentrations during and after the application of PF input alone, and compared these with their temporal evolution in response to coincident PF and CF input. The temporal evolution of unphosphorylated and phosphorylated AMPA receptors in wild-type and knockout mice in response to the two stimulation protocols is shown in Figures 4A and 4B. In wild-type mice, the application of smaller calcium pulses that are triggered by PF input alone (Figure 4A, inset) results in concentrations of unphosphorylated AMPA receptors that start exceeding their initial value after about 100 s of PF stimulation, leading to LTP induction. In *Camk2b* knockout mice, PF input without CF input rapidly decreases the level of unphosphorylated AMPA receptors, indicating the induction of LTD rather than LTP (Figure 4A). In contrast, as shown in Figure 4B, in response to the larger calcium pulses that result from the coincident activation of PF and CF input (Figure 4B, inset), wild-type mice exhibit a decrease in unphosphorylated AMPA receptors and therefore LTD, while the *Camk2b* knockout leads to an increase in the concentration of unphosphorylated AMPA receptors, which corresponds to the induction of LTP.

**Figure 4.**
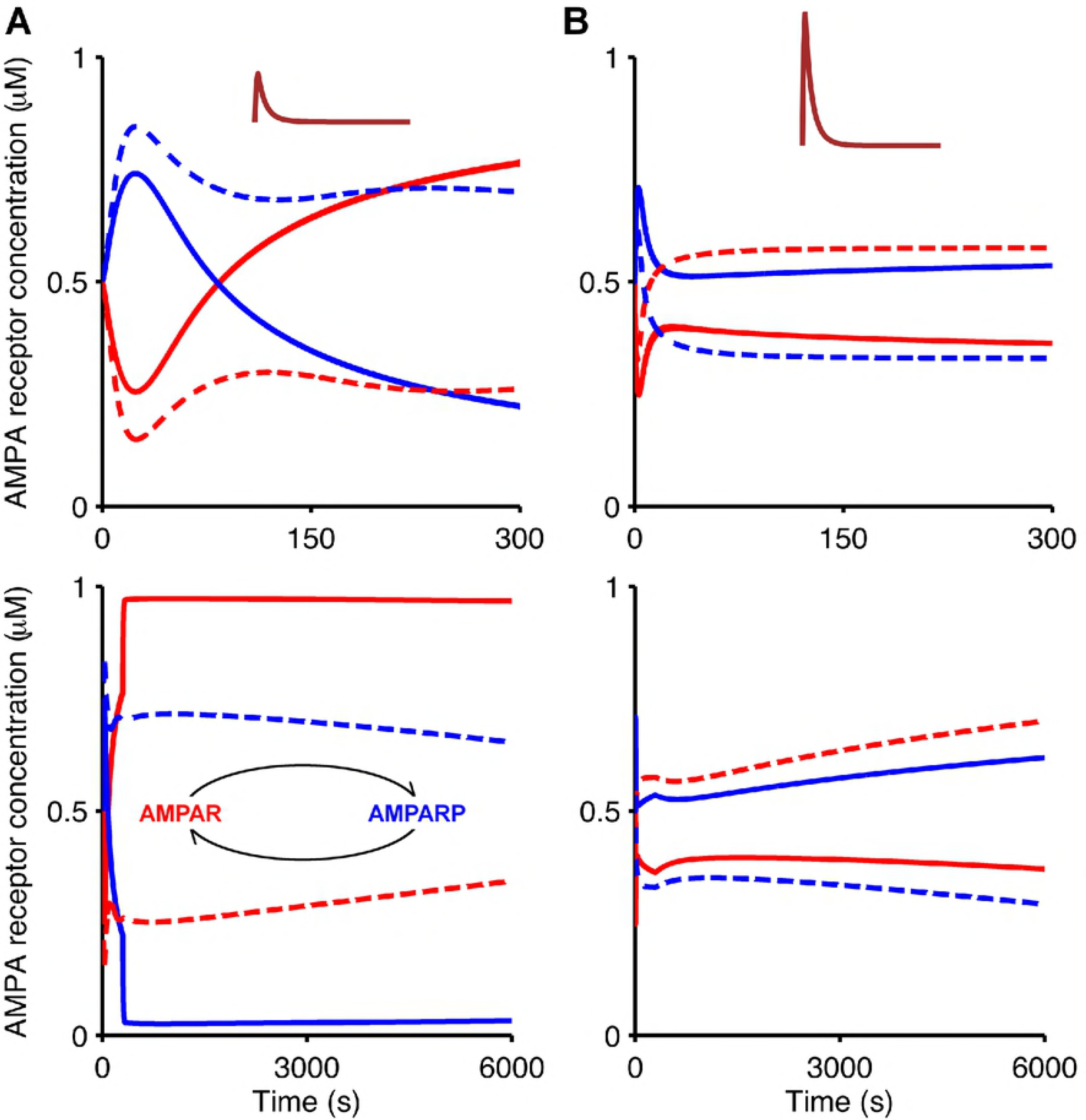
Temporal evolution of AMPA receptor phosphorylation illustrates bidirectional plasticity at PF-PC synapses in response to PF and CF input. Bidirectional plasticity at this synapse is also observed when plotting the temporal evolution of unphosphorylated and phosphorylated AMPA receptors (red and blue, respectively) for wild-type and *Camk2b* knockout mice (solid and dashed, respectively) in response to low calcium concentration resulting from the PF input alone (**A**, pulse amplitude of 1.8 *µ*M), and high calcium concentration as a result of the paired PF and CF stimulation (**B**, pulse amplitude of 10 *µ*M).

In some cases, the temporal evolution of the AMPA receptor concentrations in the simulations is biphasic. For example, PF input alone in wild-type mice leads to an initial decrease in the level of unphosphorylated receptors that is followed by an increase above the initial baseline (Figure 4A, top panel). To understand the time course of the AMPA receptor phosphorylation and dephosphorylation, we analysed the temporal evolution of the activation of CaMKII and PP2B in the presence of synaptic input (Figure 5), and their deactivation when the synaptic input ceases (Figure 6). Figures 5A and 6A indicate that the biphasic nature of the AMPA receptor phosphorylation and dephosphorylation is based on the different rates of CaMKII and PP2B activation in the model. For both wild-type and *Camk2b* knockout mice, the activation of CaMKII is faster than that of PP2B, which leads an initial increase of the number of phosphorylated (and a decrease in the number of unphosphorylated) AMPA receptors in all cases (Figure 4). However, in the presence of the F-actin binding of CaMKII in wild-type mice, the low calcium concentrations that result from PF input alone do not activate the kinase sufficiently, and the activation of PP2B surpasses the activation of CaMKII, which leads to the induction of LTP (Figures 5A and 6A, top panel). In contrast, LTD occurs for PF input alone in knockout mice, where more CaMKII is available for activation, so that the concentration of activated CaMKII exceeds that of PP2B (Figures 5A and 6A, bottom panel).

**Figure 5.**
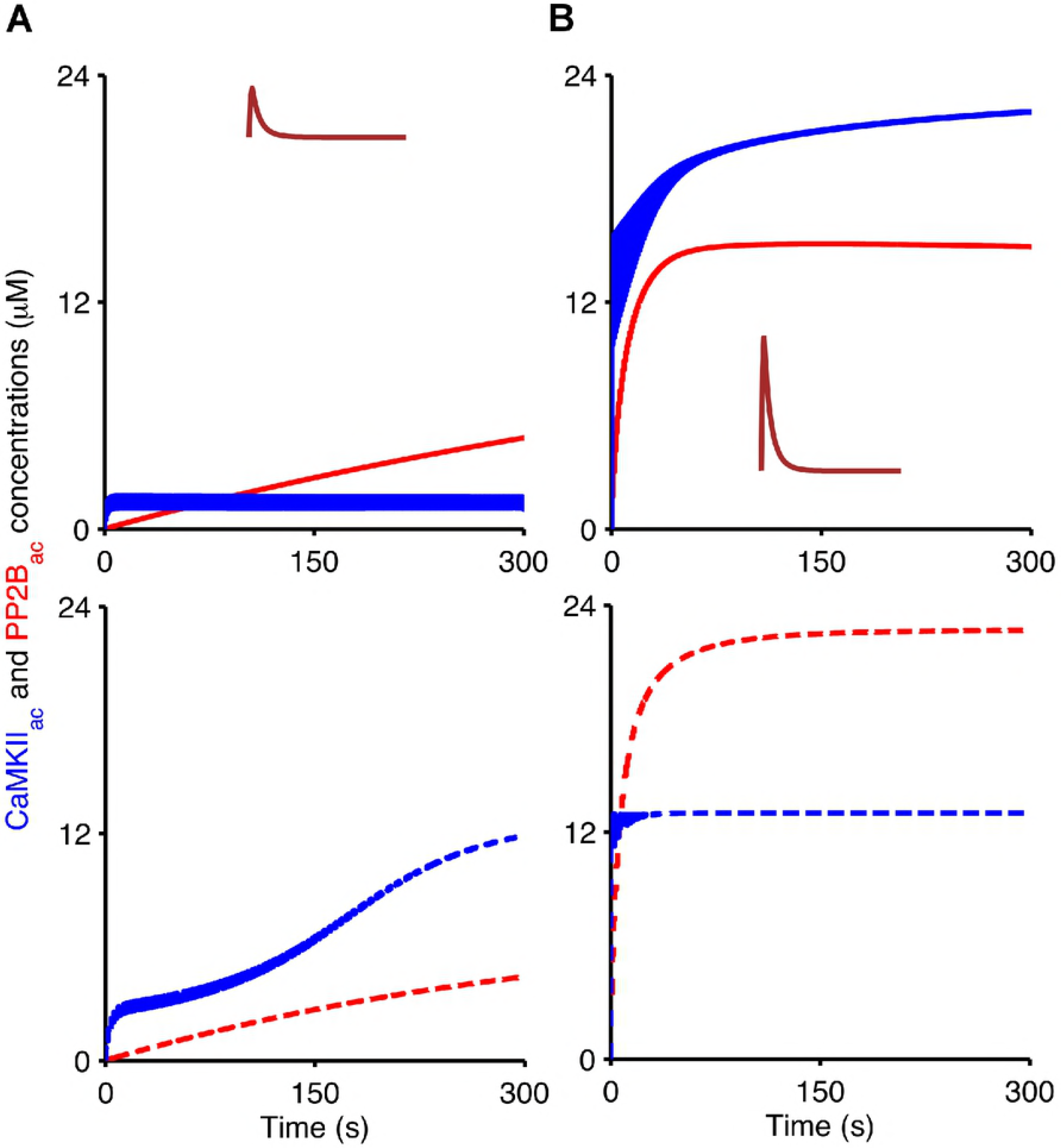
Temporal evolution of active CaMKII and PP2B during the application of PF and CF input. Evolution of concentrations of CaMKII_ac_ (blue) and PP2B_ac_ (red) in wild-type mice (top, solid) and *Camk2b* knockout mice (bottom, dashed) in response to low calcium concentrations as a result of the PF input alone (**A**, pulse amplitude of 1.8 *µ*M), and high calcium concentrations resulting from coincident PF and CF stimulation (**B**, pulse amplitude of 10 *µ*M). In this figure, simulations last 300 s which is the duration of calcium stimulation.

Figures 5B and 6B show that the opposite scenario occurs for coincident PF and CF input. The higher levels of calcium that result from the paired PF and CF stimulation result in much stronger activation of CaMKII and PP2B. In wild-type mice, the CaMKII activation exceeds the activation of PP2B, leading to LTD (Figures 5B and 6B, top panel). However, the reduction of the overall CaMKII concentration because of the *Camk2b* knockout means that the level of PP2B_ac_ in the knockout mice is higher than that of CaMKII_ac_ and LTP occurs (Figures 5B and 6B, bottom panel).

**Figure 6.**
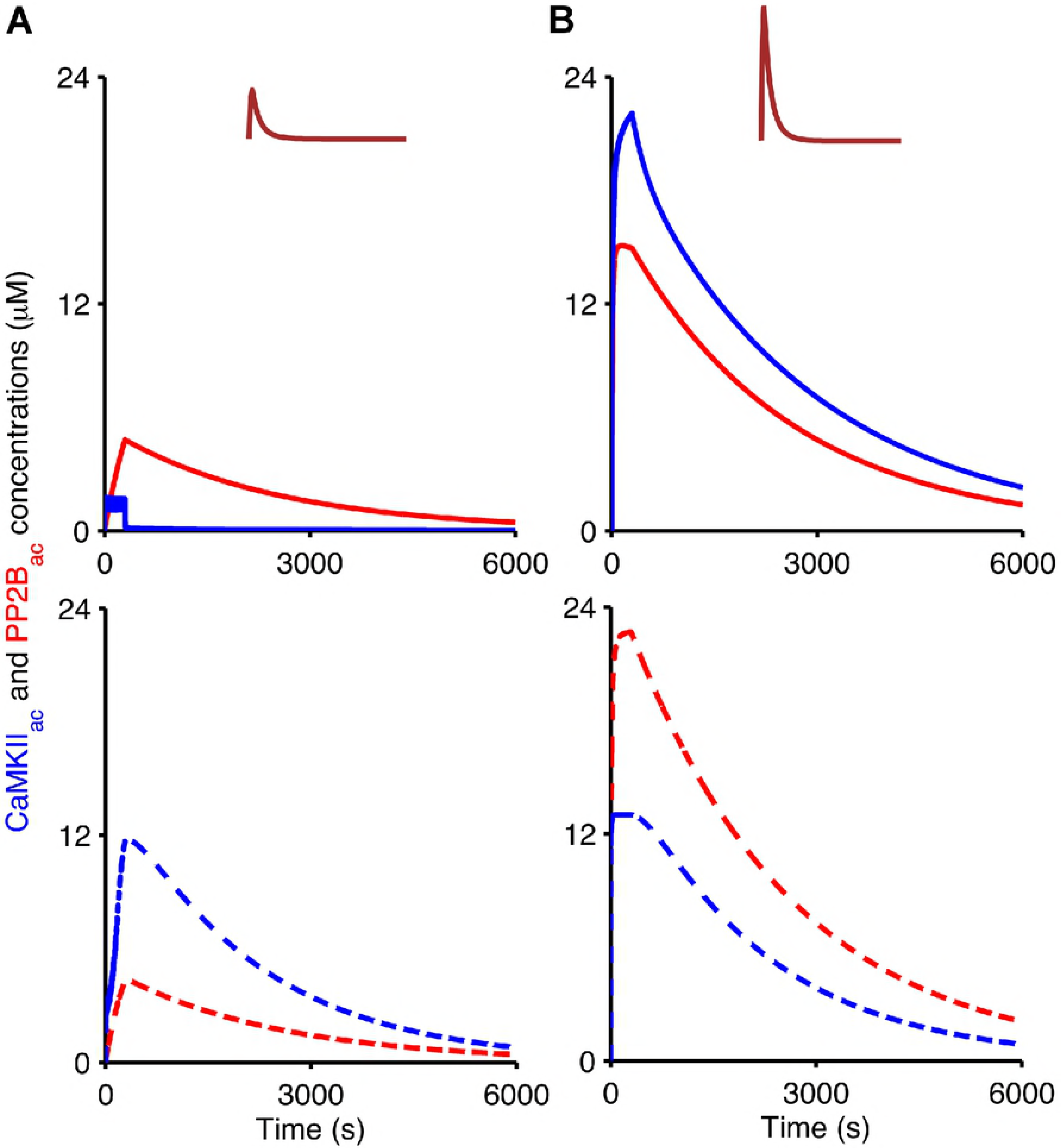
Continued temporal evolution of active CaMKII and PP2B after the application of synaptic input. Temporal evolution of CaMKII_ac_ (blue) and PP2B_ac_ (red) in wild-type mice (top, solid) and *Camk2b* knockout mice (bottom, dashed) in response to low calcium concentrations as a result of the PF input alone (**A**, pulse amplitude of 1.8 *µ*M), and high calcium concentrations resulting from coincident PF and CF stimulation (**B**, pulse amplitude of 10 *µ*M). Here, all simulations last 6000 s to investigate what occurs with CaMKII_ac_ and PP2B_ac_ concentrations after calcium input signals that last 300 s.

## Discussion

LTD of synapses between PFs and PCs in cerebellar cortex is thought to be the basis of cerebellar learning [5–7]. Postsynaptic LTD at PF-PC synapses is balanced by postsynaptic LTP, but the exact conditions and cellular mechanisms that determine the direction of synaptic plasticity induction in vivo are not fully understood. PF LTD and LTP are mediated by the phosphorylation and dephosphorylation of AMPA receptors [11, 38] and regulated by a kinase/phosphatase switch involving enzymes such as CaMKII [29, 31–33] and PP2B [14]. The control of bidirectional plasticity by CaMKII has received particular attention, after experiments by Van Woerden and collaborators with *Camk2b* knockout mice [33] indicated that the beta isoform of CaMKII plays a central role in determining the direction of synaptic weight change. In *Camk2b* knockout mice that lack *β*CaMKII, stimulation protocols that normally lead to the induction of LTD resulted in LTP, and vice versa [33]. Van Woerden et al [33] suggested that this reversal of synaptic plasticity in the *Camk2b* knockout mice could be based on binding of *β*CaMKII to F-actin in wild-type mice, which might reduce the concentration of CaMKII that is available for the phosphorylation of AMPA receptors under these normal conditions.

To test this hypothesis and investigate whether (and under which conditions) F-actin binding of CaMKII in wild-type mice, and the lack of F-actin binding in *Camk2b* knockout mice, can indeed lead to the observed reversal of synaptic plasticity at PF-PC synapses, we developed a simple mathematical model of AMPA receptor phosphorylation and dephosphorylation by CaMKII and PP2B. In our model, F-actin binding to CaMKII was included in wild-type mice that contain both *α* and *β* kinase isoforms. In contrast, in simulations of *Camk2b* knockout mice that lack *β*CaMKII the F-actin binding was omitted, and the total concentration of CaMKII was reduced to half of the value for wild-type mice. Simulation results of the model support the suggestion by Van Woerden and colleagues [33]. Moreover, they go beyond their original suggestion [33] and predict that the sign reversal of synaptic plasticity is based on a combination of three mechanisms operating at different postsynaptic calcium concentrations. At low calcium concentrations such as those that result from PF input alone, and that evoke LTP in wildtype mice, the lack of F-actin binding of CaMKII in *Camk2b* knockout mice leads to a level of active and available CaMKII that exceeds the level of active PP2B; this results in induction of LTD in these mutant mice. For the high calcium concentrations that occur in response to a classic LTD induction protocol, that is, coincident PF and CF input in wildtype mice, two mechanisms operate together. The loss of *β*CaMKII in the knockout mice causes a reduction in the total concentration of CaMKII. As a consequence, less CaMKII is available for the phosphorylation of AMPA receptors, and more Ca_4_CaM is left available for the activation of PP2B. Thus, the model predicts that the combination of a decreased kinase activity and an increased phosphatase activity is responsible for the switch from LTD to LTP for high calcium concentrations in *Camk2b* knockout mice.

Although there are many computational models that investigate the biochemical processes underlying LTD at PF-PC synapses [11, 13, 22–27], none of these models has been set up to model the induction of LTP at these synapses. Moreover, despite experimental evidence for the involvement of CaMKII in PF LTD and LTP [29, 31–33], the CaMKII pathway has, so far, not been included in the signalling cascades of existing PF-PC synaptic plasticity models. The present simple model is the first model of intracellular signalling at PF-PC synapses that focusses on the role of CaMKII and that models the induction of both LTD and LTP in wildtype and *Camk2b* knockout mice. In order to concentrate on the role of *β*CaMKII in bidirectional synaptic plasticity, the model has been kept intentionally simple. In future work, however, the biological realism of the model could be incrementally extended by adding components of existing models of PF LTD. A comparison of results obtained from our simple model with a more complex one could then be used to identify the potential contribution of other biochemical species to regulating the sign reversal of synaptic plasticity in cerebellar PCs.

Most modelling studies of intracellular signalling are based on the assumption that ionic and molecular concentrations can be described by mass action kinetics, and it is often assumed that reactions between chemical species occur in a single well-stirred compartment. As in the present model, this assumption leads to deterministic solutions of systems of ordinary differential equations. However, many signalling cascades involve very small numbers of molecules or ions, which are more faithfully represented by stochastic models [39–41]. As a complement to our deterministic model of intracellular signalling at the PF-PC synapse, a stochastic model of bidirectional plasticity at this synapse, which also accounts for spatial aspects of cellular signalling, should be developed. Results obtained from both deterministic and stochastic computational models should then be compared.

Previous models of cerebellar LTD have measured synaptic plasticity by quantifying the concentrations or densities of phosphorylated and unphosphorylated AMPA receptors [11, 24]. Indeed, this is the case for the simple model proposed here. However, Antunes and De Schutter have recently suggested that the extent of synaptic plasticity should be measured as a reduction in the number of synaptic AMPA receptors, rather than the concentration of phosphorylated receptors [42]. A next step in this research could be the inclusion of the AMPA receptor trafficking modelled by Antunes and De Schutter into the model of bidirectional plasticity at PF-PC synapses. Moreover, several concentrations of biochemical compounds and kinetic rate constants are free parameters, due to the lack of experimental data. Additional experiments will lead to an incremental improvement of the computational model, which in turn might suggest new experiments that need to be conducted.

## Supporting Information

### Bidirectional Synaptic Plasticity Model

To study the role of calcium-calmodulin dependent protein kinase II (CaMKII) in cerebellar plasticity, we first propose a rationalized version of the simple model of CaMKII activation developed by Dupont and collaborators [35]. In the model for CaMKII activation by calcium-calmodulin (Ca_4_CaM) considered here, CaMKII subunits for *Camk2b* knockout mice can be in four different states: inactive (W_i_), bound to Ca_4_CaM (W_b_), phosphorylated and bound to Ca_4_CaM (W_p_), and autonomous (W_a_): phosphorylated, but dissociated from Ca_4_CaM. For wild-type mice that contain the *α*CaMKII and *β*CaMKII isoforms, the kinase subunits can also be in states bound to filamentous actin (Ac): W_i_Ac, W_b_Ac, W_p_Ac, and W_a_Ac.

The CaMKII activation model developed by Dupont et al [35] was adapted here to include the CaMKII subunit states bound to Ac, and also to express all kinase states in concentrations rather than fractions. Therefore, the W_i_ concentration may be computed from the mass conservation relation

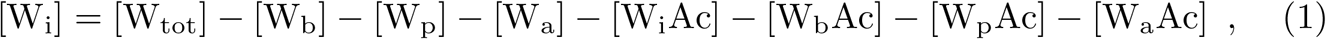

where [*x*] denotes the concentration of substance *x*, e.g. [W_tot_] is the total concentration of CaMKII.

Subunits in the W_b_, W_p_, and W_a_ states exhibit kinase activity, and can, therefore, phosphorylate CaMKII’s targets, including adjacent subunits in the CaMKII multimer. To be “ready” for phosphorylation, such adjacent subunits must be in the W_b_ state themselves. The overall phosphorylation rate associated with this process is indicated by V_a_, which is calculated using a phenomenological non-linear function of kinase subunits in the W_b_, W_p_ and W_a_ forms as in [35]

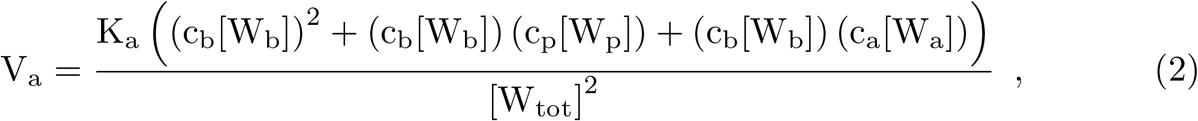

where c_b_, c_p_ and c_a_ are weighting factors proportional to the kinase activity of each active state.

The earlier model [35] includes an empirical cubic function (K_a_) to model the neighbouring autophosphorylation, allowing the mathematical model to reproduce the experimental results in [43]. The equation for K_a_ is

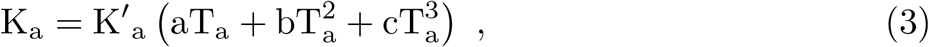

where the phenomenological rate for CaMKII autophosphorylation is K′_a_, a, b and c are parameters that Dupont et al adjusted to fit the experimental plots in [43], and T_a_ is the total fraction of active subunits represented as

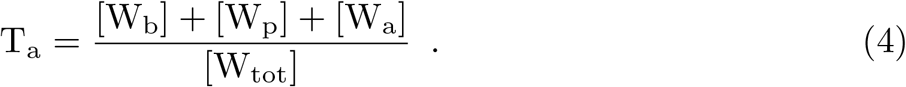

Once Ca_4_CaM binds to W_i_, the resulting W_b_ form can either be phosphorylated, release Ca_4_CaM, or bind to Ac. W_b_ is an active CaMKII subunit that can also phosphorylate AMPA receptors (AMPARs). The equation for W_b_ is therefore

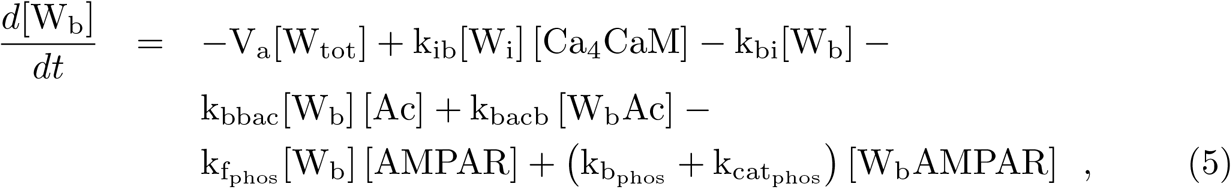

where [Ca_4_CaM] denotes the concentration of Ca_4_CaM and [W_b_AMPAR] is the concentration of W_b_ bound to AMPARs.

The W_p_ subunit can release Ca_4_CaM, switching to the W_a_ state, or bind to Ac and form W_p_Ac. As for W_b_, W_p_ also phosphorylates AMPARs. The amount of W_p_ is calculated as

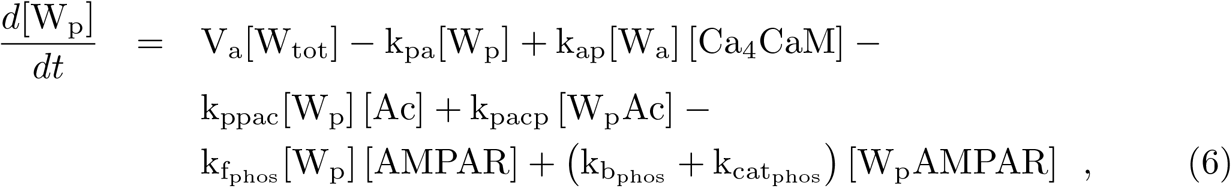

where [W_p_AMPAR] is the concentration of the complex of W_p_ bound to AMPARs.

W_a_ can bind to either Ca_4_CaM or Ac, and phosphorylate AMPARs as well. In our model, we also allow the CaMKII subunits in the W_a_ form to switch to the W_i_ state. CaMKII is therefore gradually inactivated after calcium stimulation stops. Thus,

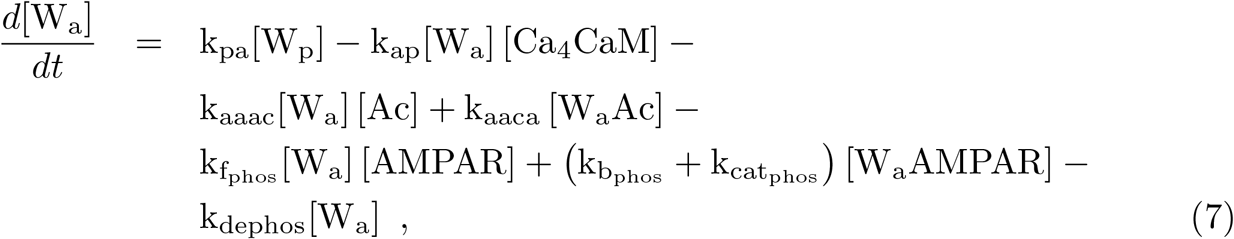

and [W_a_AMPAR] is the amount of W_a_ trapped to AMPARs.

Unlike the model in [35], the model used here does not include a separate “trapped” state in which apo-CaM (CaM without any calcium ions bound) is bound to CaMKII, mainly because dissociation of calcium and CaM cannot be distinguished experimentally (nor described thermodynamically) as two distinct processes. As a result, the values for the rate constants between the “trapped” CaM (W_t_) and W_a_ states in [35] were adopted here as k_ap_ and k_pa_.

W_i_Ac binds to Ca_4_CaM and changes to the active W_b_Ac form, or dissociates from Ac and switches to the W_i_ state. Changes in W_i_Ac concentration are

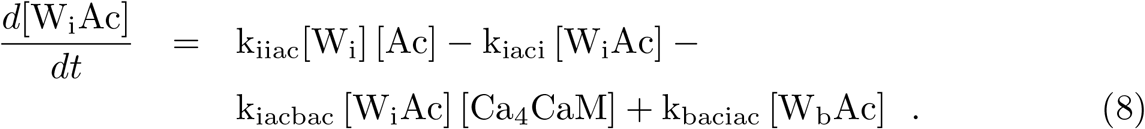

W_b_Ac can be phosphorylated by neighbouring active Ac-bound subunits: W_b_Ac itself, W_p_Ac or W_a_Ac. The autophosphorylation process for Ac-bound CaMKII is analogous to the mechanism of autophosphorylation of the kinase unbound to Ac (Equations 2, 3 and 4). The autophosphorylation rate for Ac-bound CaMKII subunits (V_ac_) is therefore

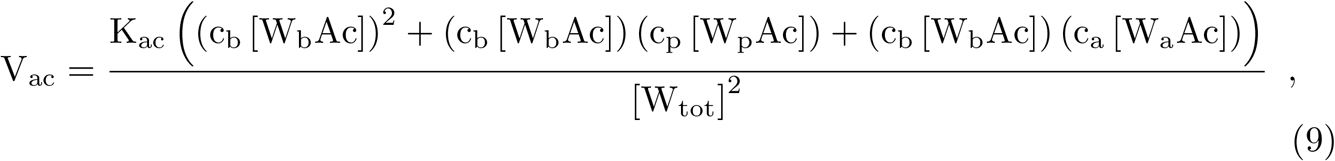

where

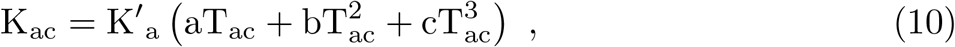

and

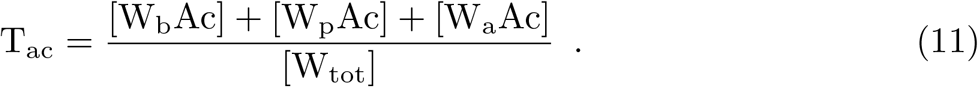

W_b_Ac can also switch to W_i_Ac once Ca_4_CaM dissociates from this kinase subunit, or dissociate from Ac and change to W_b_. Thus,

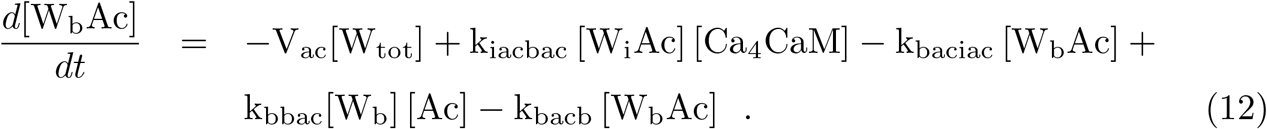

The phosphorylated Ac- and Ca_4_CaM-bound form of CaMKII can release Ca_4_CaM and switch to the W_a_Ac form, or dissociate from Ac and swap to the W_p_ state. The concentration of W_p_Ac at each time step is expressed as

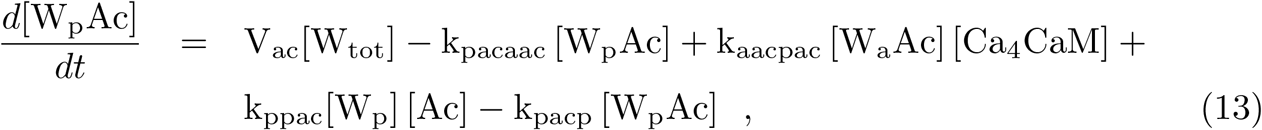

whereas W_a_Ac can bind to Ca_4_CaM and switch to W_p_Ac, or dissociate from Ac and turn into W_a_. The equation for W_a_Ac is

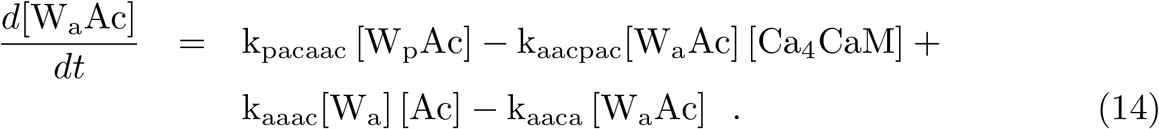

Ca_4_CaM not only activates CaMKII, but can also bind to the inactive form of PP2B (PP2B_i_). The phosphatase then gets activated and switches to the PP2B_ac_ form. PP2B_ac_ mediates the dephosphorylation of AMPA receptors. The temporal evolution of PP2B_i_ concentration is

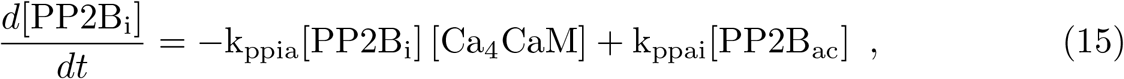

whereas PP2B_ac_ concentration changes are expressed as

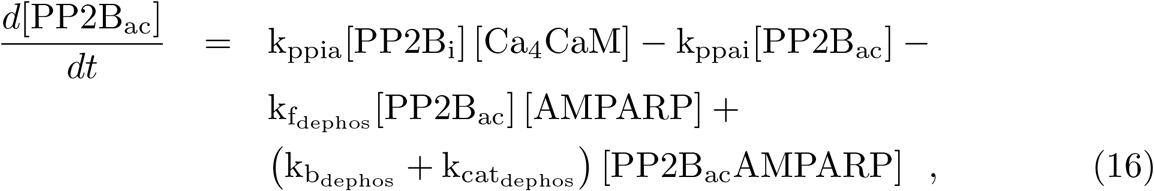

where [PP2B_ac_AMPARP] expresses the concentration of PP2B_ac_ bound to phosphorylated AMPA receptors (AMPARPs).

The evolution of unphosphorylated AMPA receptors at each time step is

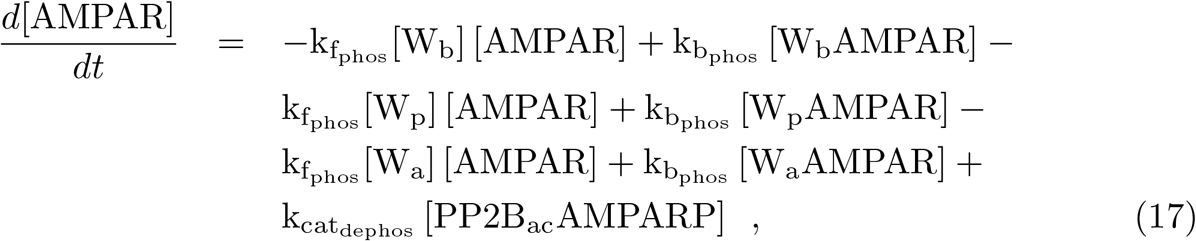

where

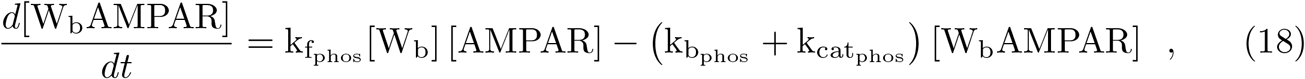

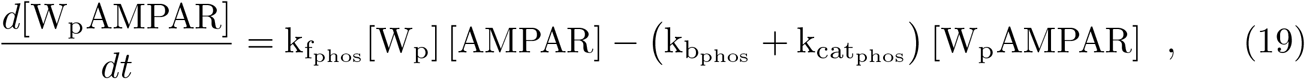

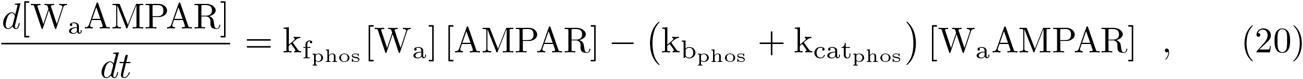

and

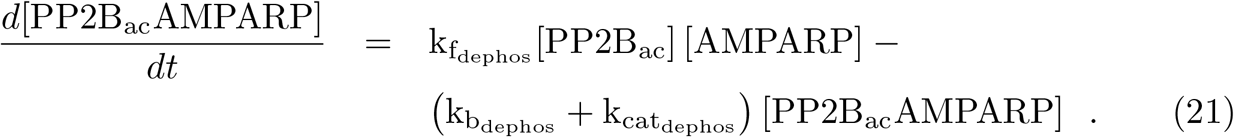

Moreover, the equation that represents the evolution of AMPARPs is

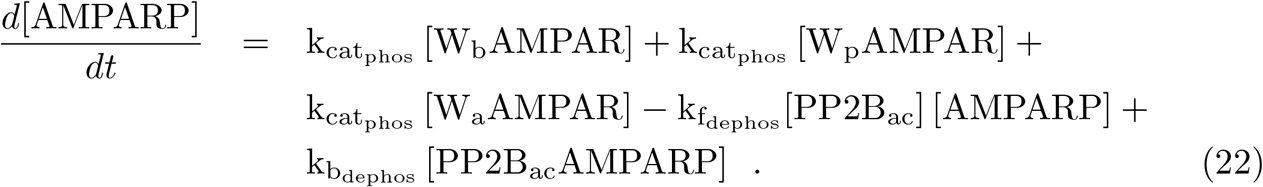

Standard protocols of long-term depression (LTD) and long-term potentiation (LTP) induction normally involve repetitive calcium stimuli (such as 1 Hz for 300 s). To generate calcium pulses with concentrations that reflect experimental data [10], our simple model that simulates the bidirectional synaptic plasticity in Purkinje cells uses a calcium dynamics model

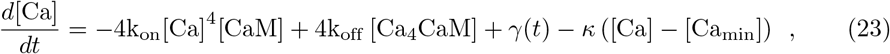

where [Ca] is the calcium concentration. The term *γ*(*t*) − *κ*([Ca] [Ca_min_]) describes the simple model of calcium dynamics we adopted, where *γ*(*t*) denotes calcium concentration increases at each time step whose values originate from an input table, *κ* is a term that reflects the calcium removal through diffusion, pumps, exchanges, and [Ca_min_] is the basal calcium concentration.

Input with high calcium influx rates (*γ*(*t*)) was used to stimulate the model to generate realistic amplitudes of calcium in response to parallel fibre (PF) alone and PF + climbing fibre (CF) stimulations (Figure 7). The temporal evolution of CaM is written as

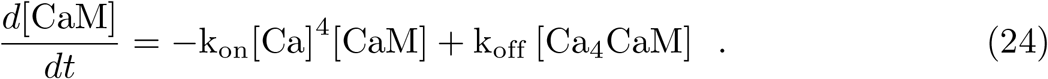

Ca_4_CaM results from the binding of four calcium ions to CaM, and activates PP2B, W_b_, W_p_, W_b_Ac and W_p_Ac. The equation that represents the evolution of Ca_4_CaM concentration is

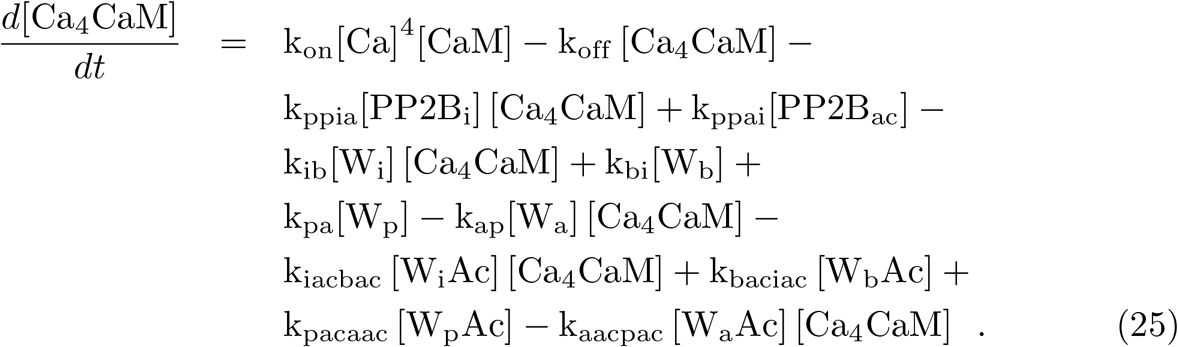

**Figure 7.**
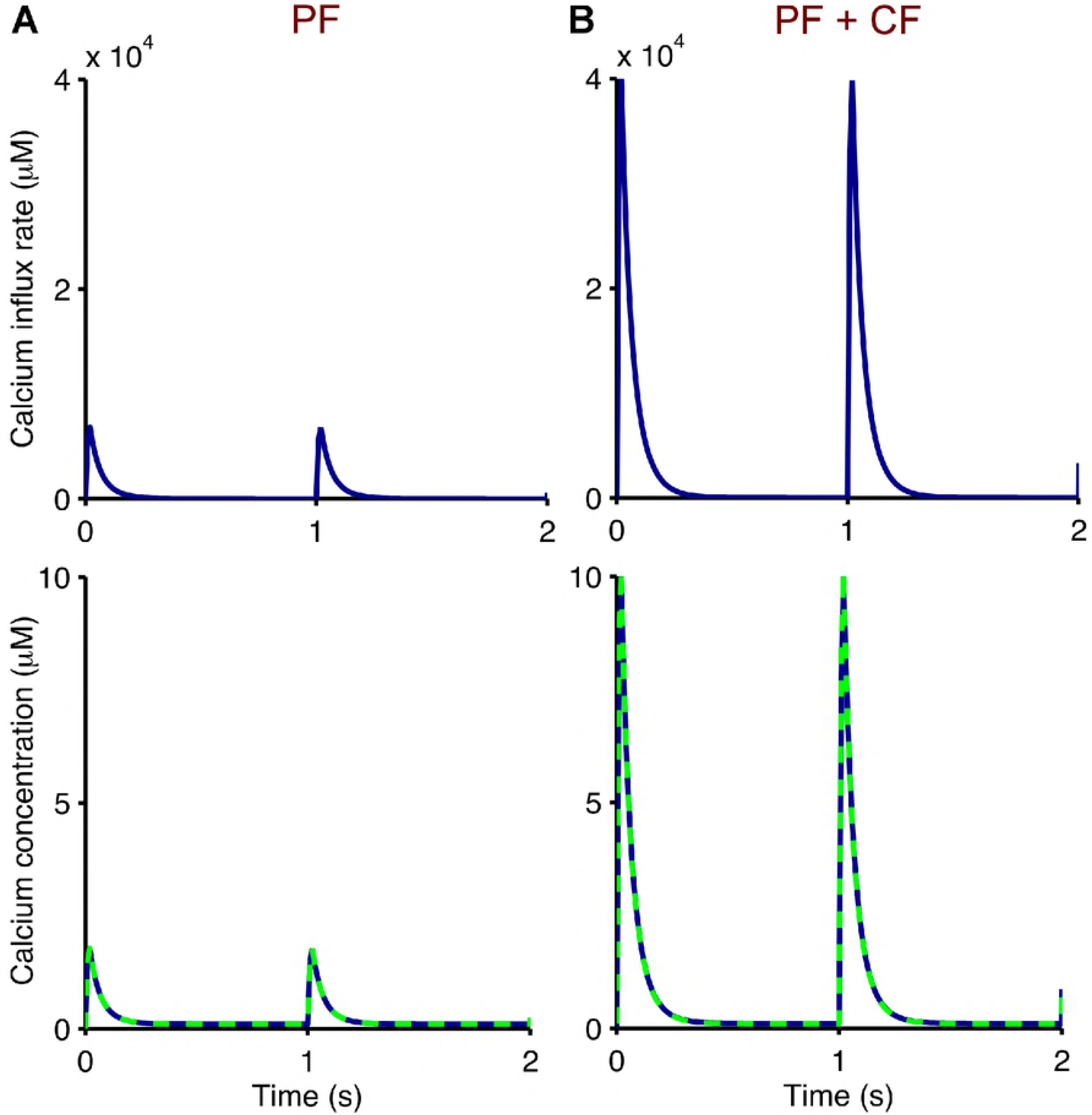
Pulsed calcium stimulation for cerebellar PF-PC synapses. Calcium influx rates (top, *γ*(*t*) in Equation 23) are used as input to our bidirectional synaptic plasticity model to generate the desired output of calcium spikes (bottom, [Ca] in Equation 23). These match experimental data in [10, 11] (dashed) which represent stimulations of PF alone (**A**, pulse amplitude of 1.8 *µ*M) and paired PF and CF (**B**, pulse amplitude of 10 *µ*M). Both calcium stimulations are applied at 1 Hz for 300 s.

**Model Parameters**

## Acknowledgments

We thank Chris De Zeeuw, Freek Hoebeek and Zhenyu Gao for helpful discussions and suggestions.

---

**Table 1.**
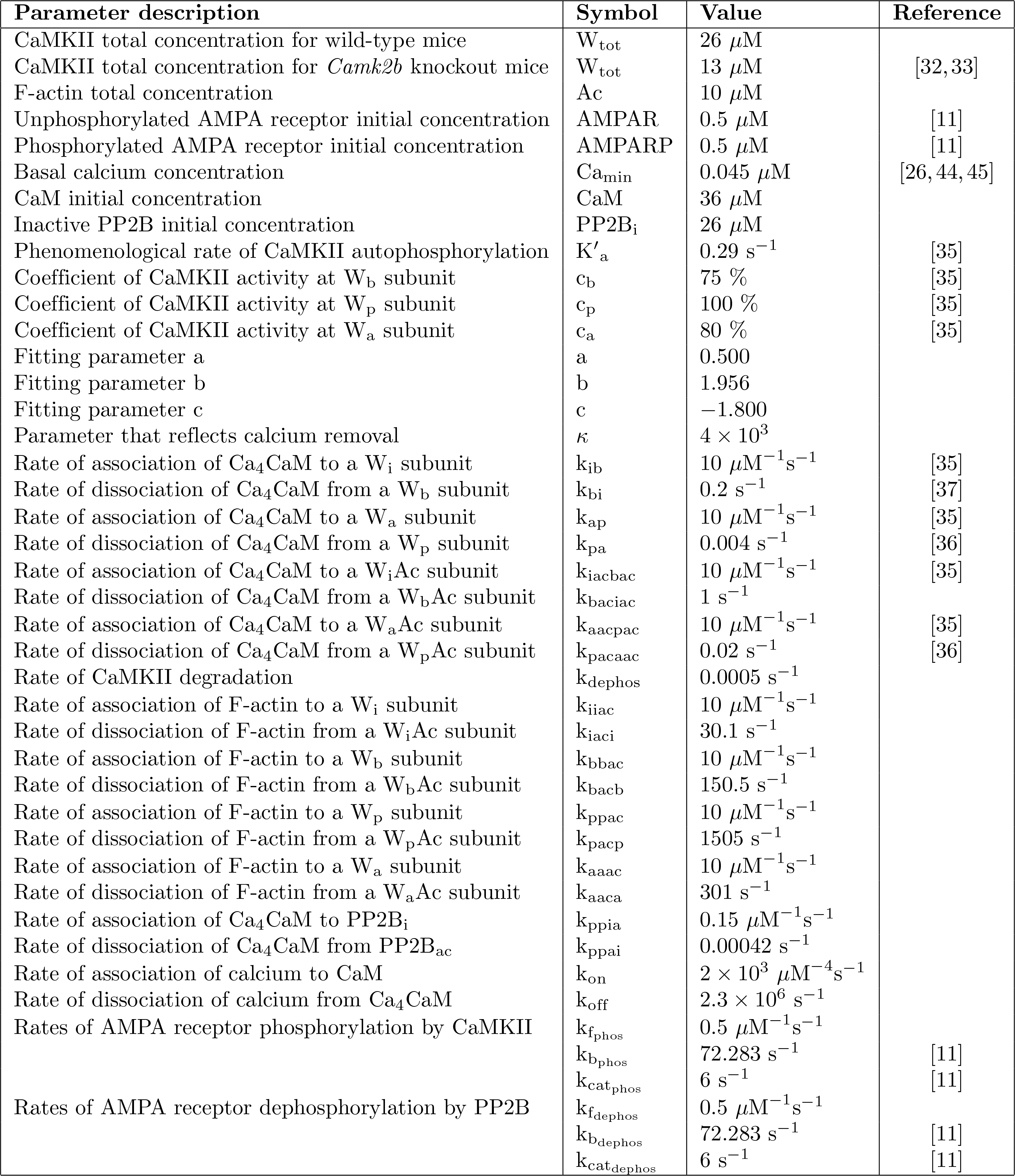
Values of kinetic parameters for modelling bidirectional plasticity at PF-PC synapses.

